# Development and validation of a gene expression score to account for tumour purity and improve prognostication in breast cancer

**DOI:** 10.1101/2024.02.23.581701

**Authors:** Marco Barreca, Matteo Dugo, Barbara Galbardi, Balázs Győrffy, NA-PHER2 consortium, NeoTRIP consortium, Pinuccia Valagussa, Daniela Besozzi, Giuseppe Viale, Giampaolo Bianchini, Luca Gianni, Maurizio Callari

## Abstract

The prevalence of malignant cells in clinical specimens, or tumour purity, is affected by both intrinsic biological factors and extrinsic sampling bias. Molecular characterization of large clinical cohorts is typically performed on bulk samples; data analysis and interpretation can be biased by tumour purity variability. Transcription-based strategies to estimate tumour purity have been proposed, but no breast cancer specific method is available yet.

We interrogated over 4400 expression profiles from 9 breast cancer datasets to develop and validate a 9-gene Breast Cancer Purity Score (BCPS). BCPS outperformed existing methods for estimating tumour content. Adjusting transcriptomic profiles using the BCPS reduce sampling bias and aid data interpretation. BCPS-estimated tumour purity improved prognostication in luminal breast cancer, correlated with pathologic complete response in on-treatment biopsies from triple-negative breast cancer patients undergoing neoadjuvant treatment and effectively stratified the risk of relapse in HER2+ residual disease post-neoadjuvant treatment.

## Background

Clinical tumours are complex ecosystems, including neoplastic, immune and endothelial cells, fibroblasts and normal epithelium. Tumour microenvironment (TME) components functionally interact one another and with the tumour cells, in a complex network of signals impacting tumour growth and progression, response to treatment and prevalence of the different cell types^1,2^.

As a consequence, tumour purity (i.e. the relative abundance of cancer cells) could represent a biologically relevant, intrinsic tumour feature. For example, a recent study in gastric cancer showed how tumour purity can predict response to chemotherapy, providing novel insights that could help improving prognostic risk stratification and facilitate treatment decision-making^3^. Similarly, association of purity with major clinical and molecular features was reported in glioma^4^. On the other hand, tissue organisation and tumour heterogeneity cause spatial variability in tumour purity. In the molecular analysis of clinical tumours, only small regions of the neoplastic lesion are typically investigated (e.g. tissue sections from a core biopsy). This introduces an extrinsic variation in tumour purity caused by the sampling procedure, which, in a pan-cancer analysis, was reported to outweigh intrinsic factors^5^.

In translational research, most high throughput experiments are performed on bulk tissue samples; consequently, variability in tumour content can influence the interpretation of molecular data and clinical decisions^6,5,7,8^. In genomic analyses, purity can impact detectability of somatic mutations^9^ and copy number alteration events^10^. In transcriptomic profiling of bulk clinical specimens, the observed profile is the result of mRNAs expressed by all different cell types present in the tumour ecosystem. A number of gene expression-based biomarkers have been developed, with some reaching clinical implementation. In breast cancer, the transcriptomics-based PAM50 molecular subtyping has been proposed and validated^11^ and is now used to aid clinical decisions^12^. Remarkably, the Normal-like subtype is allegedly considered an artifact caused by low tumour purity more than an actual cancer cell phenotype. Similarly, the classification into all other subtypes could be biased by variability in tumour purity. We and others have previously proposed the quantification of estrogen signalling and tumour proliferation as prognostic and predictive biomarkers in breast cancer^12,13^. Tumour purity could introduce also in this case a relevant bias in their quantification.

Additionally, it is becoming increasingly common to study serial samples collected during treatment^14^. Treatment is expected to have a major and patient-specific impact on tumour purity. Consequently, interpretation of differences observed between pre- and post-treatment samples could be significantly biased by changes in tumour content. At the same time, treatment-induced changes in tumour purity could represent valuable information in predicting treatment response and outcome.

Tumour purity is generally estimated by the pathologists through visual or image analysis of tumour sections and, while this method is considered the gold standard, high inter-pathologist variability and discordance have been reported in different studies^6,15^.

A few computational methods have been proposed to use molecular information for tumour purity estimation, such as gene expression^16,17^, genomic^9^ or DNA methylation profiles^9^. These methods have the potential to bypass pathology-assessment of tumour cellularity and estimate tumour purity in the very same sample used to derive the molecular profile. One of the commonly used transcriptomics-based methods is ESTIMATE^18^, which was developed pan-cancer by combining an immune and a stroma score, but not including any tumour related genes.

In this study, we exploited over 4400 samples from 9 breast cancer datasets to systematically quantify the impact of intrinsic and extrinsic factors on cellularity and to generate and validate a Breast Cancer Purity Score (BCPS). BCPS outperformed ESTIMATE in quantifying tumour purity and can be successfully used to adjust for tumour purity variability when extrinsic factors are prevalent. BCPS can also capture treatment-induced changes carrying predictive and prognostic information.

## Methods

### Dataset collection and processing

To derive and validate the BCPS, a total of 9 gene expression datasets were collected and interrogated. Information on datasets source, sample usage, sample features and data processing are detailed in Table 1.

**Table 1.**
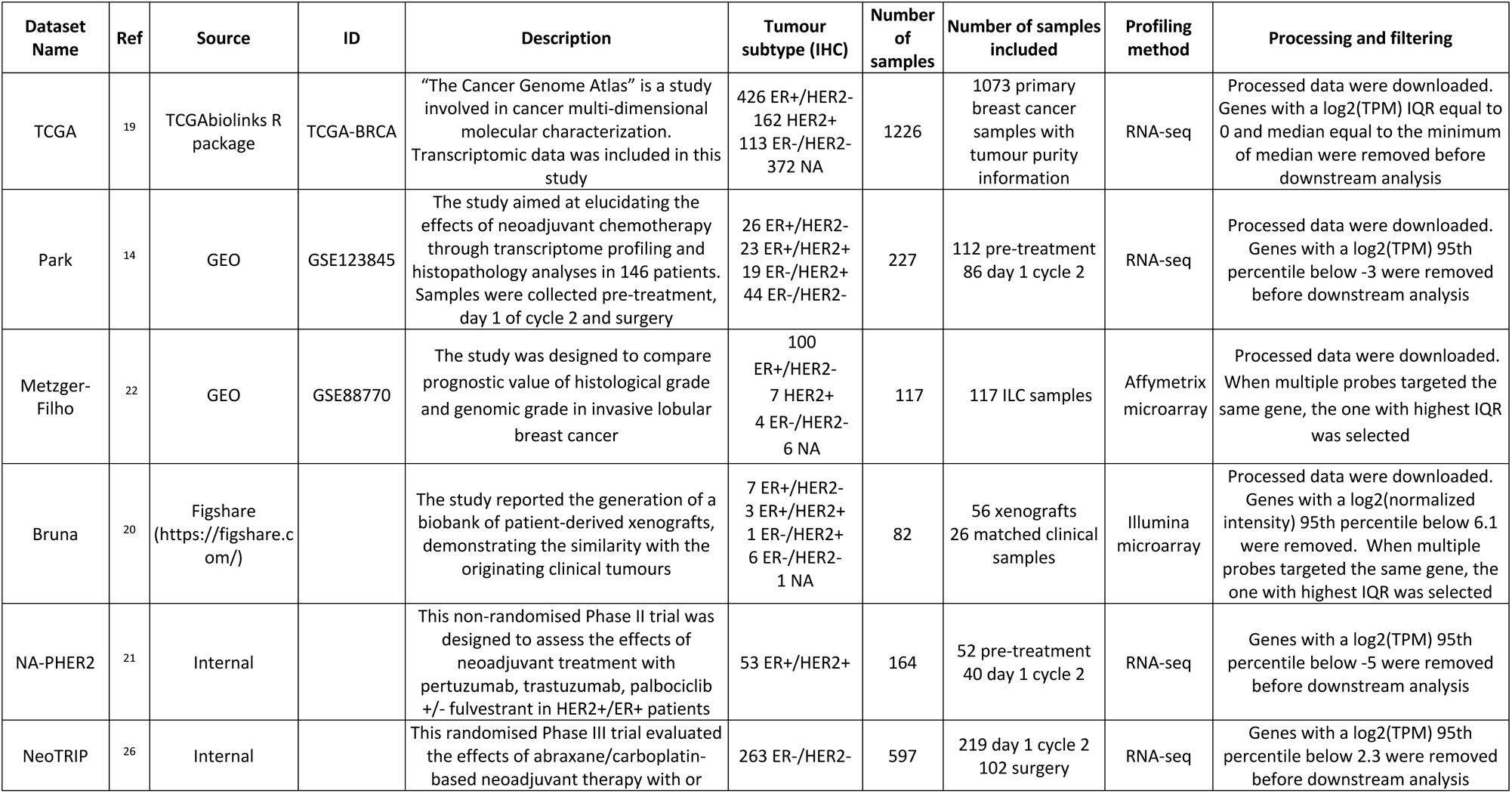
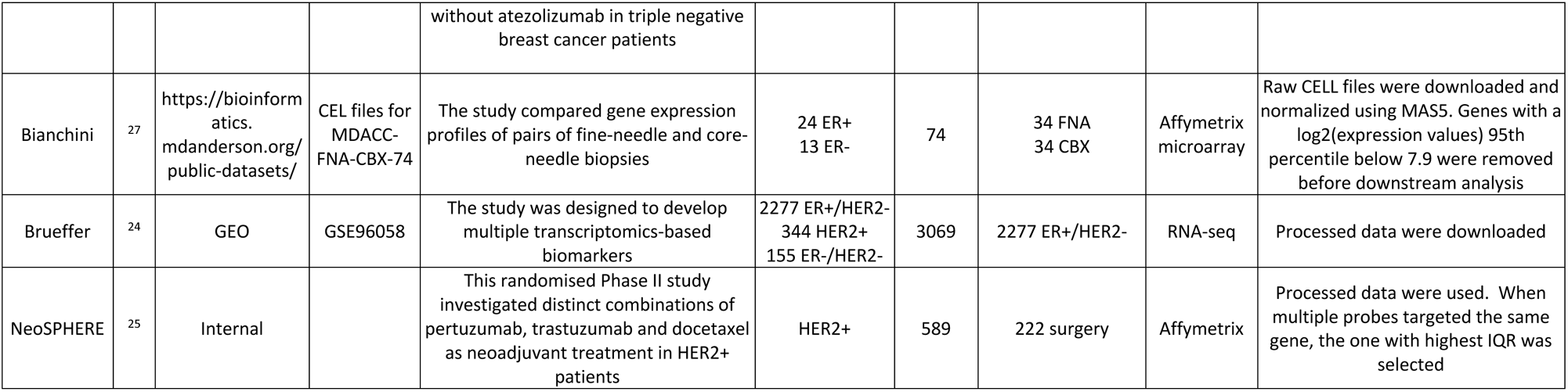
Description of gene expression datasets interrogated to derive, validate and apply the BCPS.

### BCPS identification and quantification

Genes candidate to be good reporters of tumour cellularity were identified according to the scheme in Figure 2. From the intersection of candidate genes obtained from each of the four interrogated datasets (i.e. TCGA^19^, PDX^20^, NA-PHER2^21^, Metzger-Filho^22^), 5 genes consistently associated with tumour content and 4 consistently associated with stroma content were selected. The two genesets were used together to perform a directional single-sample geneset enrichment analysis as implemented in the *singscore* (version 1.14.0) R package^23^. Higher score indicates higher tumour purity.

### ESTIMATE score computation

ESTIMATE score^18^ was computed using the *estimate* (version 1.0.13) R package. The score is computed by combining a stromal and an immune score. To improve readability and comparability with the BCPS, ESTIMATE score was multiplied by −1 to have higher values indicating higher tumour purity.

### Statistical analyses

#### Variable associations

Spearman’s correlation was calculated to evaluate the association between two continuous variables. Differences between correlation values obtained on the same set of data were compared using *cocor* (version 1.1.4) R package. Student’s t-test (*stats* R package, version 4.1.2) and ANOVA analysis (*car* R package, version 3.0.14) were calculated to evaluate the statistical significance of association of a continuous variable with two or multiple classes, respectively.

#### Variance Component Analysis

Variance component analysis (VCA) was computed to evaluate the impact of clinico-pathological features on tumour purity as implemented in the *VCA* (version 1.4.3) R package. Samples with missing values in any of the variables were excluded.

#### ROC curve analysis

To evaluate the ability of a variable to discriminate between two classes, Receiver Operating Characteristic curve (or ROC curve) and associated area under the ROC curve (AUC) were computed, as implemented in the *ROCR* (version 1.1.11) R package. Differences in AUC values were statistically evaluated using *pROC* (version 1.18.0).

#### Differential expression analysis

Differential expression analysis was performed using the *limma* (version 3.50.0) R package. Nominal p values were corrected for multiple testing using the Benjamini-Hochberg method. Genes with False Discovery Rate (FDR)<0.05 and absolute log fold change >1 were considered significantly differentially expressed, unless otherwise specified.

#### Survival analysis

To evaluate the association with survival for single or multiple variables, univariate or multivariate Cox regression analysis as implemented in the *survival* (version 3.2.15) R package was performed. In the TCGA and Brueffer^24^ datasets, overall survival was the clinical endpoint. In the NeoSPHERE^25^ dataset, distant event-free survival (DEFS) was considered. To quantify and compare the prediction power of distinct models, the concordance index, as implemented by *stat* (version 4.1.3) R package, and the 7-years area under the ROC curve (AUC), as implemented by *survivalROC* (version 1.0.3) R package, were computed.

## Results

### Association of pathology-assessed cellularity with clinico-pathological features and patient’s outcome

To obtain insights into the relative impact of intrinsic and extrinsic factors on breast cancer sample tumour content, we investigated whether pathology-assessed tumour cellularity is associated with biologically and clinically relevant features in breast cancer. Associations were evaluated in the TCGA, Metzger-Filho, Park and NA-PHER2 datasets, where tumour purity quantification by expert pathologists was available (Table 1). We found multiple significant associations as reported in Figure 1A. In the TCGA, the invasive lobular cancers (ILC) had lower purity than other subtypes (Figure 1A and Figure S1A). At the same time, in the Metzger-Filho dataset, including only ILCs, cellularity was significantly different between distinct ILC subtypes, with the lobular Classic subtype showing the lowest cellularity (Figure S3B). The proliferation marker Ki67 had a weak correlation with cellularity in the Metzger-Filho dataset (ρ = 0.19, p = 0.047), but the same association was not confirmed in Park and NA-PHER2 datasets (Figure 1A, Figure S2M, Figure S3C and Figure S4B). On the contrary, high grade was consistently associated with higher cellularity (Figure 1A, Figure S2A, Figure S3A). Breast cancer subtypes, either defined by the combination of ER and HER2 status or according to PAM50 classification, showed a significant association with cellularity in both TCGA and Park datasets. In particular, triple-negative or basal-like tumours had, on average, the highest cellularity (Figure 1A, Figure S1E-F and Figure S2E-F). Finally, in NA-PHER2 samples, stromal tumour-infiltrating lymphocytes (sTILs), but not intraepithelial tumour-infiltrating lymphocytes, were negatively correlated with tumour cellularity (ρ = −0.33, p = 0.018) (Figure 1A and Figure S4C-D).

**Figure 1.**
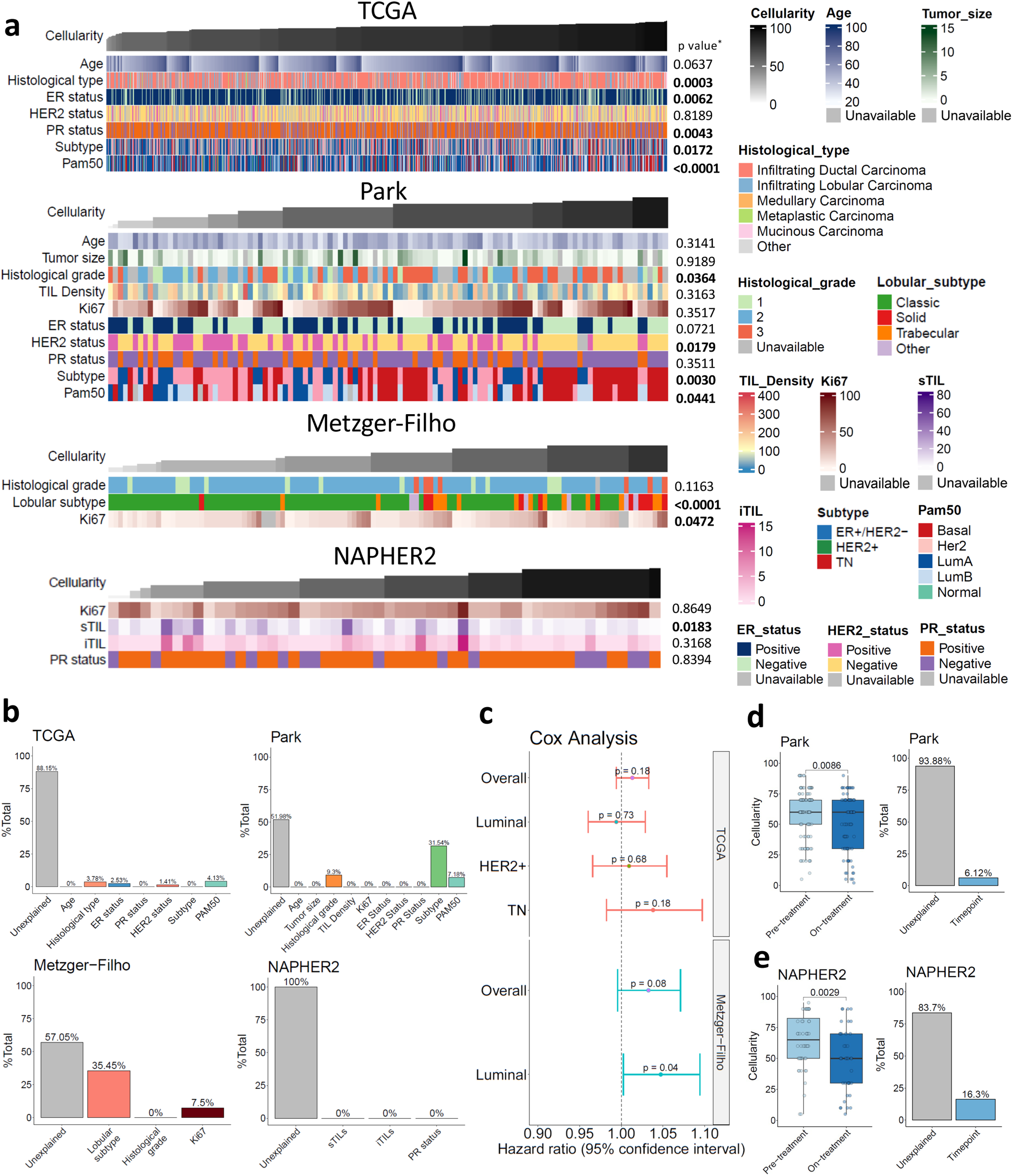
Association of pathologist’s estimated tumour purity with molecular and clinico-pathological variables in breast cancer. **a)** Landscape of association of available molecular and clinico-pathological variables with cellularity in four breast cancer datasets: TCGA (n = 1073), Metzger-Filho (n = 117), Park (n = 112) and NA-PHER2 (n = 52) pre-treatment samples. (*\** association between purity and continuous variables was assessed by Spearman’s correlation, association with two categorical groups was assessed by Student’s *t* test, and association with multiple categorical groups was evaluated by one-way ANOVA). **b)** Variance component analysis (VCA) for each dataset computed for samples with no missing information (TCGA, n = 690; Park, n = 72; Metzger-Filho, n = 111; NA-PHER2, n = 52). The analysis estimated the proportion of total variance explained by the provided variables. **c)** Forest plot of Cox regression univariate analysis evaluating association of cellularity with overall survival in TCGA (n = 1073) and Metzger-Filho (n = 117) datasets. Samples were evaluated overall and stratified by subtype (TCGA: 426 Luminal, 162 HER2+, 113 TN; Metzger-Filho: 100 Luminal). **d)** Cellularity changes in on-treatment biopsy (n = 86) compared to pre-treatment (n = 112) in the Park dataset. The impact of the timepoint on tumour purity was evaluated by Student’s *t* test and VCA. **f)** Same analysis as in (d) for the NA-PHER2 dataset (n = 52, pre-treatment biopsy; n = 40, on-treatment biopsy).

To quantify the contribution of each clinico-pathological variable to the overall tumour cellularity variance, we performed a variance component analysis (VCA) for each dataset (Figure 1B). In line with the analysis above, the histological type and grade, together with the molecular subtypes, explained the highest percentage of total variance. However, 52-100% of cellularity variance in each dataset was not explained by the included variables.

Next, we evaluated whether tumour cellularity was associated with distant metastasis free survival in the TCGA and Metzger-Filho datasets, overall and for each subtype defined by ER and HER2 status (Figure 1C). Cellularity was significantly associated with prognosis only in lobular luminal cases of the Metzger-Filho dataset (p = 0.04).

Finally, in the Park and NA-PHER2 datasets, transcriptomic profiles were obtained from pre- and on-treatment biopsies. Because of treatment-induced tumour cell death, an overall reduction in tumour cellularity could be expected and was observed (p = 0.009 and p = 0.003, respectively). The biopsy timepoint explained 6.12% of the variance in the Park dataset and 16.3% in the NA-PHER2 dataset (Figure 1D-E).

In summary, this analysis denoted that, in breast cancer, intrinsic tumour biology factors can affect tumour cellularity. However, over half of the variability observed in clinical specimens undergoing molecular characterization was not explained by the main clinico-pathological features and could be related to tumour sampling.

### Development of a Breast Cancer Purity Score (BCPS)

Cellularity estimation by the pathologist is not always available and might not refer to the same tumour region undergoing molecular characterization. Consequently, we aimed at identifying a gene expression signature able to estimate tumour purity in a bulk transcriptomic analysis of clinical breast cancer samples. As detailed hereafter, to generate the BCPS we interrogated four distinct datasets: Bruna, NA-PHER2 (pre-treatment samples), NeoTRIP^26^ (surgical samples) and Metzger-Filho (Figure 2 and Table 1).

**Figure 2.**
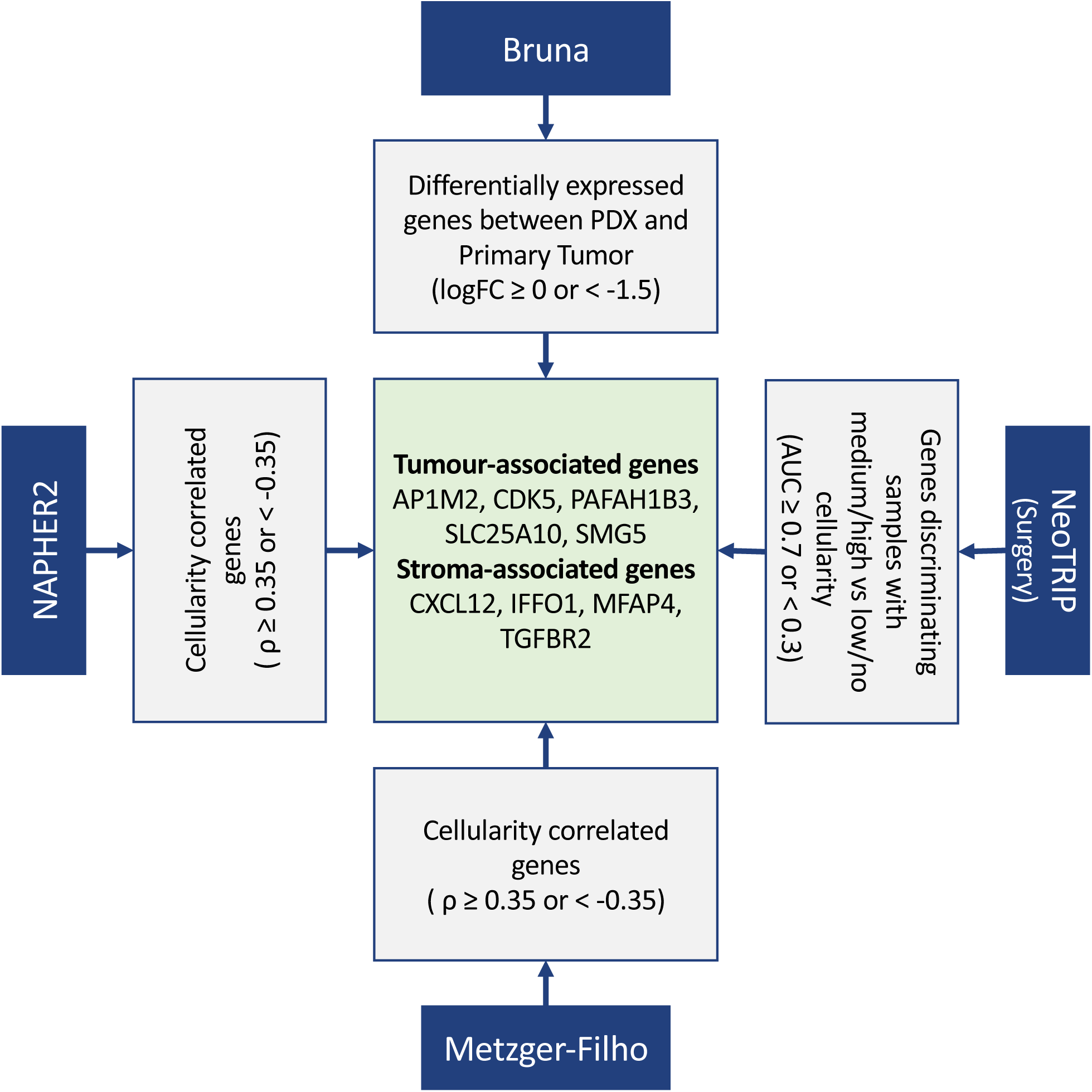
BCPS generation. Workflow involving four distinct datasets leading to the definition of the BCPS. In the NA-PHER2 and Metzger-Filho datasets, the correlation between tumour purity and the expression values of each gene was computed. In the Bruna dataset, primary tumours were compared to matched patient-derived xenografts to identify candidate tumour-specific and stroma-specific genes, exploiting the loss of human stroma during engraftment. In the NeoTRIP dataset, the ROC curve AUC was estimated for each gene considering surgical samples with medium/high or low/no cellularity, as annotated by expert pathologists. By applying for each analysis the indicated thresholds, 5 tumour-associated genes and 4 TME-associated genes were selected to generate the BCPS.

We correlated gene expression with tumour cellularity in the NA-PHER2 (n = 52) dataset, identifying 733 genes with a positive correlation with tumour cellularity (ρ > 0.35) and 1009 genes with a negative correlation (ρ < −0.35). The same analysis was performed in the Metzger-Filho (n = 117) dataset, identifying 49 genes with a positive (ρ > 0.35) and 248 genes with a negative (ρ < −0.35) correlation with cellularity. Hence, we considered surgical samples from the NeoTRIP dataset grouped into two categories based on expert pathologist evaluation: high/mid tumour content or low/no tumour content. We assessed the ability of each gene to distinguish between the two classes by ROC curve analysis, identifying 570 genes with AUC ≥ 0.7 and 330 genes with AUC < 0.3.

As a complementary strategy, in the Bruna dataset, transcriptomic profiles of patient-derived xenografts (PDXs) were compared with matched clinical samples from which the xenografts originated. Since the human stroma is completely lost during engraftment and replaced by mouse stroma^20^, genes expressed by the human tumour microenvironment are expected to be downregulated in such comparison. At the same time, tumour specific genes are expected to be similarly expressed or upregulated in PDXs. The fold changes for selected genesets, expected to be tumour specific or stroma-specific, are reported in Figure S5A-C as proof of concept. In total, 236 stroma-specific and 5679 tumour specific genes were identified, candidate to be good reporter of tumour content.

From the candidate genes selected in each dataset (Figure S5D-G), we derived a consensus list of 5 tumour-associated (AP1M2, CDK5, PAFAH1B3, SLC25A10, SMG5) and 4 stroma-associated (CXCL12, IFFO1, MFAP4, TGFBR2) genes (Figure 2). This set of genes were used for a single-sample geneset enrichment analysis providing the BCPS, proportional to sample tumour purity, as characterised in the following section.

### BCPS evaluation of performance and comparison with ESTIMATE score

To evaluate the ability of the BCPS to estimate tumour sample purity and to compare its performance with the commonly used ESTIMATE score^18^, we interrogated four additional independent datasets: TCGA, Park, NeoTRIP (on-treatment samples) and Bianchini^27^ (Figure 3 and Table 1).

**Figure 3.**
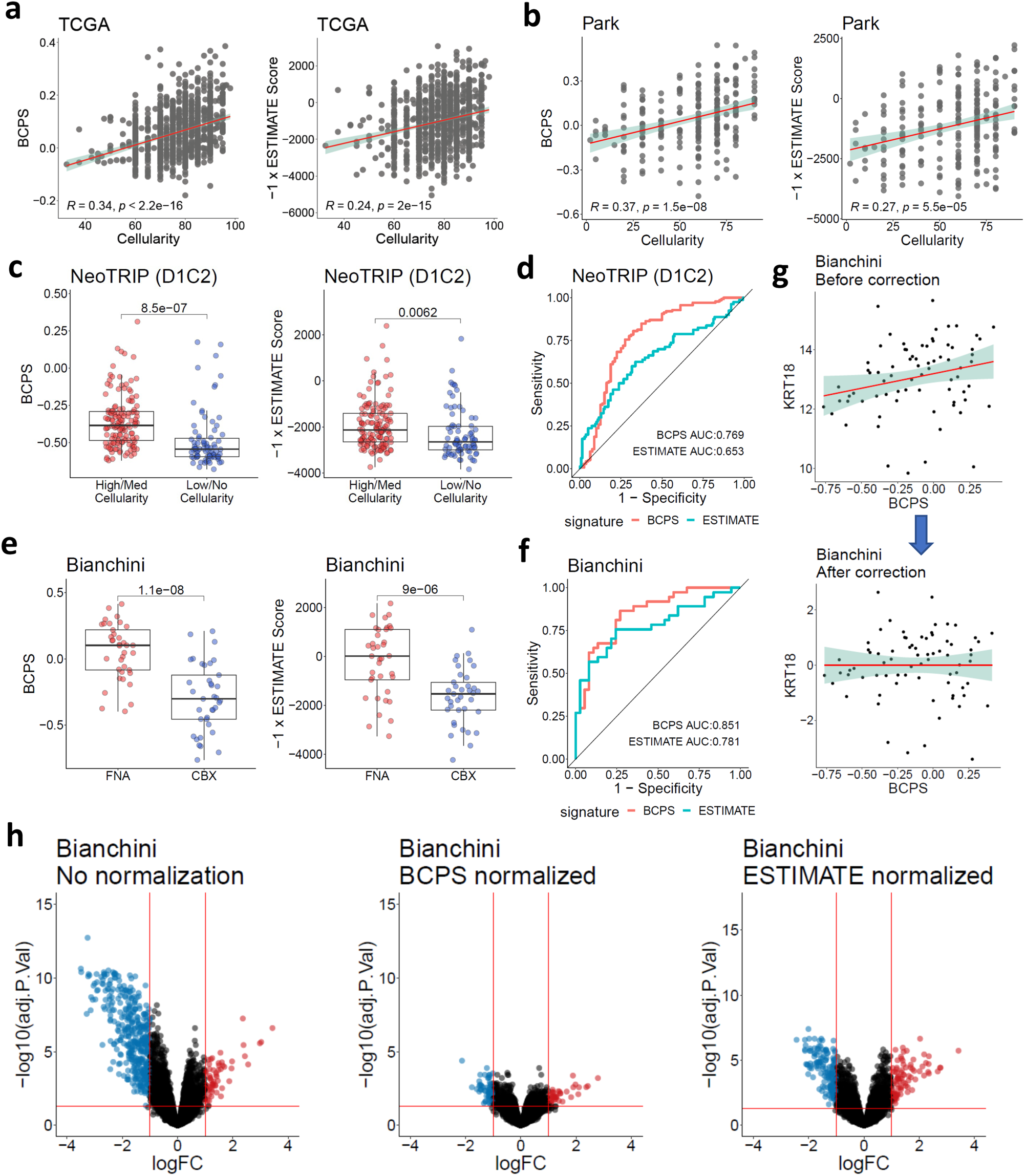
BCPS validation and comparison with ESTIMATE score. Evaluation of the BCPS and comparison with ESTIMATE score in the TCGA, Park, NeoTRIP and Bianchini datasets. **a)** Spearman’s correlation between the pathologist-estimated cellularity and either ESTIMATE or BCPS in TCGA (n = 1073). **b)** Same analysis as in (a) for the Park dataset (n = 225). **c)** ESTIMATE score and BCPS values measured in samples with high/medium tumour content or low/no tumour content in the NeoTRIP dataset (n = 219, on-treatment biopsy); two-sided Student’s *t* test. **d)** ESTIMATE score and BCPS ability to discriminate between the two classes in (c) quantified by AUC. **e)** ESTIMATE score and BCPS values measured in core biopsies (CBX) and matched fine-needle aspirations (FNA, n = 37 pairs) from the Bianchini dataset; two-sided Student’s *t* test. **f)** ESTIMATE Score and BCPS ability to discriminate between the two classes in (e) quantified by AUC. **g)** Example of KRT18 gene expression correction using the BCPS and linear regression to remove the impact of tumour purity. The Bianchini dataset was used. **h)** Volcano plots of differential gene expression analysis between FNA and CBX samples of the Bianchini dataset. The analysis was performed without any correction and after normalising gene expression using the BCPS or ESTIMATE scores.

In the TCGA and Park datasets, BCPS had a significantly higher correlation with cellularity than ESTIMATE (ρ = 0.34-0.37 vs ρ = 0.23-0.27 respectively; correlation difference p < 0.001) (Figure 3A-B). ESTIMATE was strongly inversely correlated with a gene expression-based immune score^28^ (ρ = - 0.87 and ρ = −0.90 in TCGA and Park, respectively), while cellularity and BCPS showed weaker correlations (Figure S6).

The association with tumour content was also evaluated in the on-treatment biopsies of the NeoTRIP dataset (Figure 3C-D). Samples were grouped into high/medium tumour content or low/no tumour content, as established by expert pathologists, and the ability of the two scores to discriminate between these two classes was quantified by Student’s t-test and ROC curve analysis. BCPS better discriminated between the two groups, achieving a significantly higher AUC (BCPS AUC = 0.77, ESTIMATE AUC = 0.65, p < 0.001).

We then evaluated the BCPS and ESTIMATE score in the Bianchini dataset. This dataset contains matched clinical samples obtained by either fine-needle aspiration (FNA) or core-biopsy (CBX). The first sampling procedure is known to enrich for tumour cells while CBX better preserves the stromal content. Indeed, both BCPS and ESTIMATE score were higher in the FNA samples (paired Student’s t-test p = 1.1×10^−8^ and p = 9×10^−6^, respectively). ROC curve analysis highlighted a significantly higher AUC for the BCPS compared to ESTIMATE score (BCPS AUC = 0.85, ESTIMATE AUC = 0.78, p = 0.046) (Figure 3E-F).

In the Bianchini dataset, differences between matched FNA and CBX are expected to be primarily related to sampling differences affecting tumour content. Consequently, it represents a relevant setting where to quantify such bias and evaluate the ability of a purity score to correct for it. To this aim, we performed a paired differential analysis between FNA and CBX introducing either no correction or normalising the data using the BCPS or ESTIMATE score. BCPS-normalized data were obtained by taking the residuals of the linear regression models evaluating the relationship between BCPS and each gene. An example of gene expression before and after correction is shown in Figure 3G. Differential analysis between FNA and CBX before correction identified 60 up-regulated genes and 409 down-regulated genes. Data correction using the ESTIMATE score reduced the number of differentially expressed genes to 89 up-regulated and 160 down-regulated, but only 40 up-regulated and 65 downregulated genes were observed in data corrected using the BCPS, further confirming the higher performance of the BCPS in estimating breast cancer content in clinical samples (Figure 3H). This analysis indicated that in all situations where extrinsic factors are expected to largely overweigh intrinsic factors in affecting tumour purity, BCPS is a useful and effective tool to take tumour purity into consideration and correct for it.

### BCPS recapitulates cellularity associations with clinico-pathological factors

For the TCGA and Park datasets, not used for the BCPS development, we evaluated the association with available clinico-pathological factors, as reported for the pathologist cellularity in Figure 1.

BCPS was significantly associated with the same variables that were significantly associated with cellularity in both the TCGA and Park datasets (Figure 4A and Figure 1A). VCA analysis was quantitatively different but qualitatively similar (Figure 4B and Figure 1B), and the BCPS was significantly lower in on-treatment compared to pre-treatment samples in the Park dataset, as observed for the pathologist’s cellularity (Figure 4D and Figure 1D). Finally, the BCPS was not significantly associated with survival in any subtype in the TCGA dataset (Figure 4C and Figure 1C). These results support the validity of the BCPS in quantifying tumour purity and identifying intrinsic factors affecting it.

**Figure 4.**
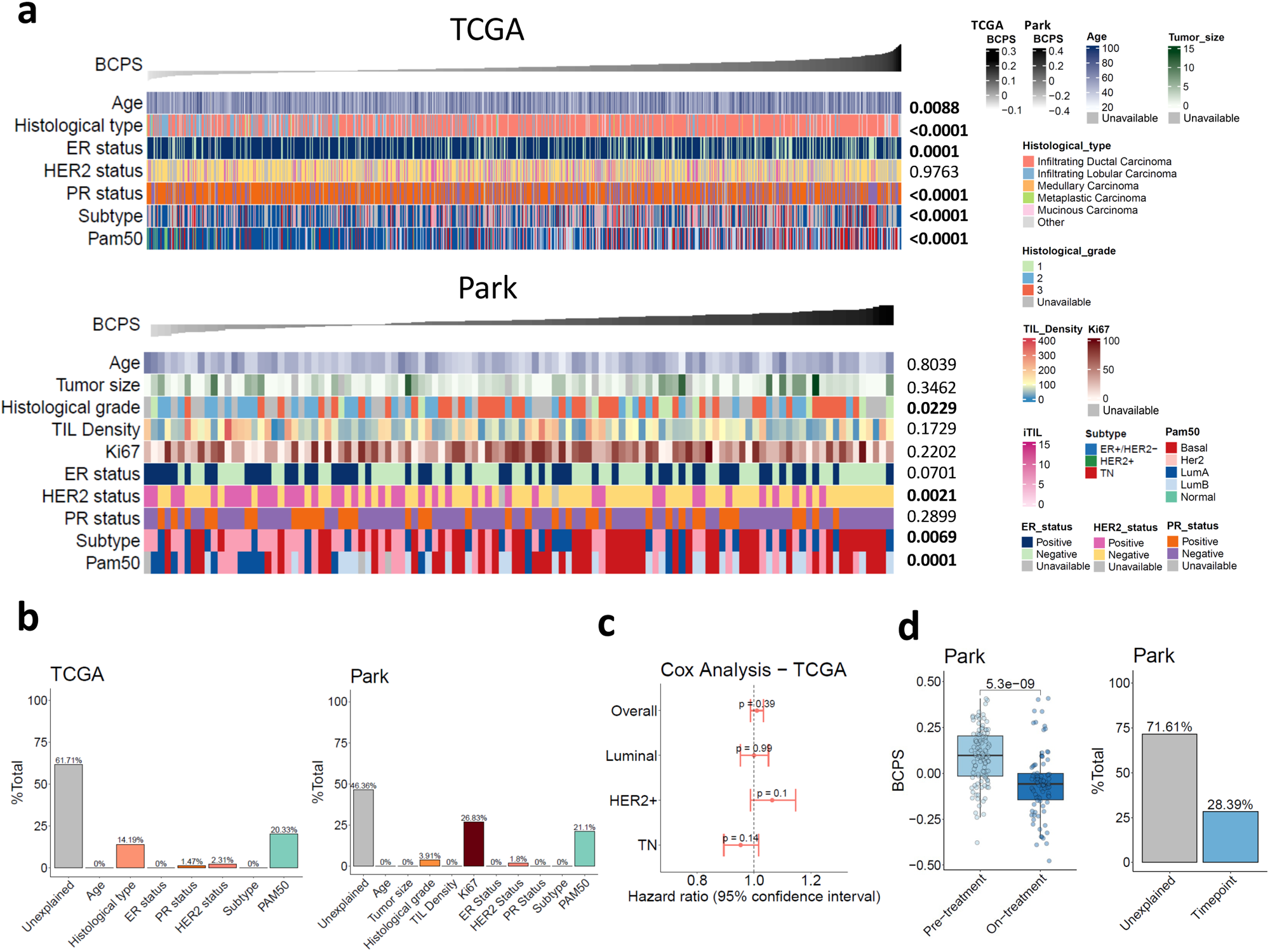
Association of the BCPS with clinico-pathological factors in the TCGA and Park datasets. **a)** Landscape of association of available molecular and clinico-pathological variables with cellularity in the TCGA (n = 1073) and Park (n = 112) datasets. (*\** association between purity and continuous variables was assessed by Spearman’s correlation, association with two categorical groups was assessed by Student’s *t* test, and association with multiple categorical groups was evaluated by one-way ANOVA). **b)** Variance component analysis (VCA) for each dataset computed for samples with no missing information (TCGA, n = 690; Park, n = 72). The analysis estimated the proportion of total variance explained by the provided variables. **c)** Forest plot of Cox regression univariate analysis evaluating cellularity association with overall survival in the TCGA (n = 1082) dataset. Samples were evaluated overall and stratified by subtype (426 Luminal, 162 HER2+, 113 TN). **d)** Cellularity changes in on-treatment biopsy compared to pre-treatment in the Park dataset (T1 = 112, T2 = 86). The impact of the timepoint on tumour purity was evaluated by Student’s *t* test and VCA.

### Use of the BCPS for prediction of prognosis and response to treatment

A typical goal in transcriptomic data analysis of clinical samples is the identification of genes or signatures associated with specific clinical endpoints. Quantification of expression could be affected by tumour content^7^. We previously developed and validated an ER-related and a proliferation-related metagenes as predictors of long-term outcome in ER+/HER2-breast cancer^13^. Here we applied the metagenes to 2277 ER+/HER2-breast cancer samples from the Brueffer dataset (Table 1). A multivariate Cox model with interactions, including the ER- and proliferation metagenes, and the BCPS explained better the survival data than a bivariate model without the BCPS (likelihood ratio test p = 0.035). The improvement was confirmed by a higher c-index and higher 7-years AUC when the BCPS was included in the model (Figure 5A-B).

**Figure 5.**
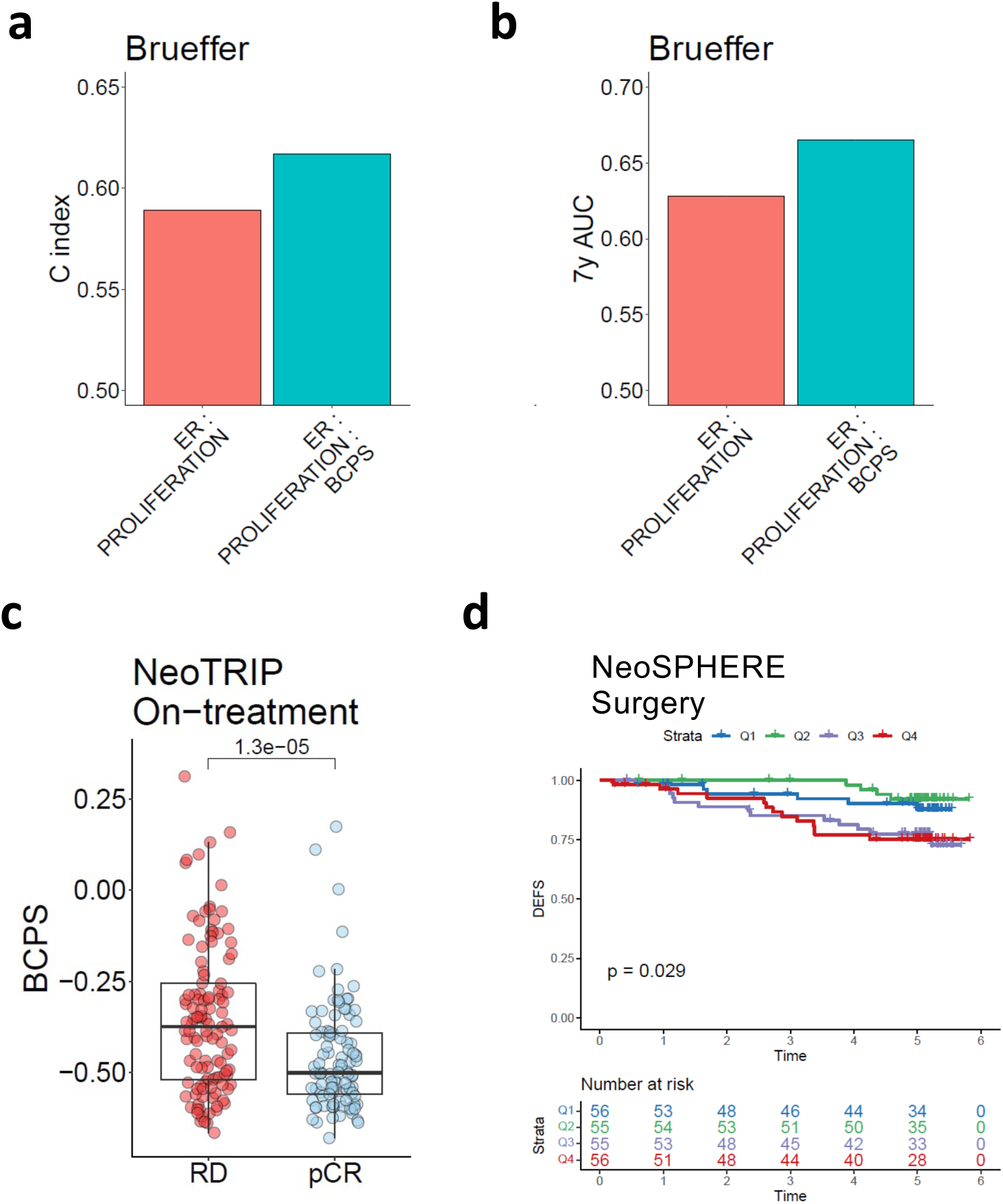
Use of the BCPS to improve prognostication in breast cancer. **a-b)** In ER+/HER2-samples from the Brueffer dataset (n = 2277) 7-years overall survival was predicted using a multivariate Cox model with interactions including an ER and a Proliferation metagene with or without the BCPS. C-index (a) and 7-years AUC (b) were computed for the two models highlighting a performance improvement when the BCPS was included. **c)** Association of the BCPS with pCR in on-treatment biopsies from the NeoTRIP dataset (n = 54). **d)** BCPS quantified in the surgical samples of the NeoSPHERE trial was associated with DEFS. Four groups based on the BCPS quartiles were identified and represented by Kaplan-Meier curves and differences evaluated by log-rank test.

Next, we focused on BCPS values in on-treatment biopsies of the NeoTRIP dataset, where the BCPS was significantly lower (p = 1.3×10^−3^) in cases eventually achieving pCR (Figure 5C). This indicates that, despite a possible sampling bias, evaluation of cellularity in on-treatment biopsies could help early prediction of patients responding or not to neoadjuvant treatments.

A third example where evaluation of the BCPS could provide valuable information is in surgical samples obtained after neoadjuvant treatment. Cellularity was included as one of the factors determining the Residual Cancer Burden, which has been associated with long term survival^29^. Here we evaluated the BCPS in post-treatment surgical samples from the neoadjuvant clinical trial NeoSPHERE^25^. The BCPS significantly stratified long-term patients’ risk by quantifying the amount of residual tumour left (Figure 5D).

## Discussion

Multiple cell types constitute the cancer ecosystem and the prevalence of each cell type can be influenced by intrinsic and extrinsic factors. Prevalence of neoplastic cells, or tumour cellularity, is typically estimated by the pathologist on histological tissue sections. In four distinct breast cancer datasets we found significant associations between pathologist-estimated cellularity and clinico-pathological features. We observed the highest cellularity in Grade 3 and ER-negative breast cancers, and the lowest in low grade luminal A tumours. Additionally, cellularity was affected by the histological type, higher in Invasive Ductal Carcinoma compared to Lobular but also variable between distinct Lobular subtypes. A weak negative correlation between cellularity and sTILs was observed in the NA-PHER2 trial. Overall, this confirmed that intrinsic factors have an impact on tumour cellularity in breast cancer, in line with what was previously reported in other cancer types^3,4^. However, VCA analysis indicated that more than half of the variance is not explained by the investigated factors, supporting the idea that extrinsic factors, including sampling bias, can affect specimen cellularity.

Bulk transcriptomic data analysis and interpretation is directly affected by the tumour content in the specimen^7^. Depending on the specific context and aims of the analysis, if extrinsic factors are expected to prevail, it is advantageous to consider tumour cellularity in data modelling.

Tumour cellularity evaluated by the pathologist is not always available and a low concordance has been reported between pathologist estimation and estimation based on DNA or RNA profiling data^6^. This could be partially explained by spatial variability between the sample evaluated by the pathologist and the sample or sections undergoing nucleic acid extraction and quantification. Moreover, the pathologist quantification is partially subjective and qualitative, with a significant interobserver variability being previously reported^6,15^.

Transcriptomics-based approaches to estimate tumour purity have been proposed. The most used is the ESTIMATE score by Yoshihara and colleagues^18^. It merges a stroma and immune signature to estimate the relative abundance of neoplastic cells and TME. It was developed to have a pan-cancer validity, but it is overly related to immune infiltration and does not include tumour specific genes. Quon and colleagues developed ISOPure^30^, a gene expression method using normal and tumour profiles to correct the latter for normal contamination. ISOPure substantially assumes that the TME would resemble the originating normal tissue, but it is well documented that this is not the case, with stroma, immune and endothelial cells dramatically altering their phenotypes as a result of the interplay with neoplastic cells^1^.

The BCPS was developed by interrogating large sample cohorts to identify the best reporter genes in a context-specific data-driven way. We covered all breast cancer subtypes, aiming at including tumour specific genes not primarily affected by the tumour subtype. We also included data obtained by different platforms; consequently, we believe that the selected consensus genes will perform well independently of the technology used. In four independent datasets we could validate the performance of the BCPS as a reporter of tumour purity. The BCPS systematically outperformed the ESTIMATE score^18^. It had significantly higher correlation with pathologist’s cellularity than ESTIMATE, significantly better identified samples with low/no tumour cells and better controlled for the sampling bias in matched CBX and FNA samples.

In the TCGA dataset, we did reproduce ESTIMATE correlation values reported by Aran et al.^5^, and the BCPS correlation with the pathologist cellularity was the highest when considering the other purity estimation metrics reported in the study (i.e. ABSOLUTE^9^ and LUMP^31^ based on genomics and methylation data, respectively). Correlation with the pathologist estimation was still moderate, but in line with previous reports^6,31^ for the reasons outlined above.

The BCPS is a simple and easy to compute score, proportional to the quantity of neoplastic cells in a clinical breast cancer specimen. As a reporter of tumour purity, the BCPS could be informative to study intrinsic factors affecting the tumour content, but it can also be used to control for tumour purity in the analysis and interpretation of bulk transcriptomic data. We showed two strategies to include the BCPS in the analyses. The first is to directly adjust genes or signatures for the BCPS-estimated effect of tumour purity on their expression levels. This strategy was effective in remarkably reducing the bias between matched CBX and FNA samples, with FNA known to be enriched in tumour cells compared to CBX. Of note, after adjusting for tumour purity, the expression levels of tumour specific genes or signatures are interpretable as the expression levels in the tumour compartment. However, TME marker genes and signatures, e.g. immune-related, are often interpreted as a proxy for a specific TME cell type prevalence. In this case, adjusting for tumour content could lead to capture cell density more than their absolute quantity, possibly providing complementary information.

A second strategy is to include the BCPS in data modelling, for example in the fitting of survival models. In this context, the other included variables could be, to some extent, biologically related with tumour purity. This was the case using ER-related and proliferation-related metagenes to predict survival in ER+HER2-breast cancer. ER expression is associated with tumour purity, either estimated by the pathologist or using the BCPS. Similarly, the higher tumour purity in Grade 3 and triple-negative tumours as well as the higher tumour purity in luminal B compared to luminal A tumours, suggests a biological link between proliferation and tumour content. In this case, including the BCPS as a covariate in the model was advantageous compared to a direct BCPS adjustment of the metagenes.

Finally, the BCPS has potential as prognostic or predictive biomarker in specific contexts. In the NeoTRIP neoadjuvant trial, BCPS-estimated low tumour content in on-treatment biopsies was associated with the achievement of pCR at the end of treatment. We also found in the Neosphere trial that the BCPS-estimated cellularity after neoadjuvant treatment of HER2+ breast cancer was associated with long-term outcome, mimicking the prognostic role of pathologist-estimated cellularity, which is one of the key components of the highly prognostic Residual Cancer Burden^32^.

## Conclusions

In this study, we developed and validated a straightforward tool to estimate tumour content from bulk transcriptomic breast cancer data, useful to explore the role of tumour purity, aid data interpretation and improve prognostication. The framework presented here could be successfully applied to other cancer types.

## List of abbreviations

ANOVA: ANalysis Of VAriance
AUC: Area under the ROC curve
BCPS: Breast Cancer Purity Score
CBX: Core-Biopsy
DEFS: Distant Event-Free Survival
EFS: Event-Free Survival
FDR: False Discovery Rate
FFPE: Formalin-Fixed, Paraffin-Embedded
FNA: Fine-Needle Aspiration
FPKM: Fragments Per Kilobase Million
IDC: Invasive Ductal Carcinoma
ILC: Invasive Lobular Cancer
iTIL: intraepithelial Tumour-Infiltrating Lymphocyte
OS: Overall Survival
pCR: pathological Complete Response
PDX: Patient-derived xenograft
ROC: Receiver Operating Characteristic
sTIL: stromal Tumour-Infiltrating Lymphocyte
TME: Tumour Microenvironment
VCA: Variance Component Analysis

## Declarations

### Ethics approval and consent to participate

All ethical approvals have been obtained as previously reported^21,33^.

### Consent for publication

Not applicable

### Availability of data and materials

Reference code to implement and use the BCPS is available under Academic Free License v. 3.0 at: https://github.com/BarrecaMarco/BCPS. Repositories and IDs of publicly available datasets used in this study are reported in Table 1. Gene AUC values computed in the NeoTRIP surgical samples and used for the BCPS development are available in Supplementary_File_1. BCPS values in NAPHER2, NeoTRIP and NeoSPHERE datasets are also reported in Supplementary file 1.

### Authors’ contributions

MC conceived the study and planned the computational and statistical analyses; MB performed the computational and statistical analyses; MD, BG and DB contributed to the computational analysis; PV curated the clinical information for the internal datasets; GV evaluated the tumour content in the internal datasets; MC and MB drafted the manuscript; GB and LG contributed to study design and clinical interpretation of the results. The Na-PHER2 consortium and the NeoTRIP consortium ran the respective clinical studies collecting the samples and clinical information. All authors contributed to subsequent drafts and agreed with submission of the manuscript for publication.

## NA-PHER2 Consortium

Luca Gianni, Fondazione Michelangelo

Giancarlo Bisagni, AUSL – IRCCS, Reggio Emilia, Italy

Marco Colleoni, IEO, European Institute of Oncology, IRCCS, Milan, Italy

Lucia Del Mastro, IRCCS Ospedale Policlinico San Martino - University of Genova, Genova, Italy

Claudio Zamagni, IRCCS Azienda Ospedaliero-Universitaria Bologna, Italy

Mauro Mansutti, Udine Academic Hospital, Udine, Italy

Milvia Zambetti, San Raffaele Hospital, Milano

Antonio Frassoldati, Azienda Ospedaliero-Universitaria S. Anna, Ferrara, Italy

## NeoTRIP Consortium

Luca Gianni, Fondazione Michelangelo, Milano

Filippo Montemurro, Fondazione Piemontese per la Ricerca sul Cancro, Candiolo

Claudio Zamagni, Policlinico S. Orsola Malpighi SSD Oncologia Medica Addarii, Bologn

Lucia Del Mastro, Università degli Studi di Genova, Ospedale Policlinico San Martino, Genova

Carmelo Bengala, Istituto Toscano Tumori, Grosseto

Marco Colleoni, Istituto Europeo di Oncologia, Milano

Gabriella Mariani, Fondazione IRCCS Istituto Nazionale Tumori, Milano

Anna Gambaro, Ospedale Luigi Sacco, Milano

Stefania Zambelli, Ospedale San Raffaele, Milano

Giampaolo Bianchini, Ospedale San Raffaele, Milano

Giancarlo Bisagni, AUSL IRCCS Arcispedale S. Maria Nuova, Reggio Emilia

Stefania Russo, Ospedale S. Maria della Misericordia, Udine

Chiun-Sheng Huang, National Coordinator, National Taiwan University Hospital, Taipei

Shou-Tung Chen, Changhua Christian Hospital, Changhua City

Ming Feng Hou, Kaohsiung Medical University Chung-Ho Memorial Hospital Cancer Center, Kaohsiung City

Liang-Chih Liu, China Medical University Hospital, Taichung City

Ling Ming Tseng, Taipei Veterans General Hospital, Taipei

Catherine Kelly, National Coordinator, Mater Misericordiae University Hospital, Dublin

Seamus O’Reilly, Cork University Hospital, Cork

Patrick Morris, Beaumont Hospital, Dublin

John Kennedy, St. James’s Hospital, Dublin

Miriam O’Connor, University Hospital Waterford, Waterford

Richard Greil, National Coordinator, IIIrd Medical Department with Hematology, Medical Oncology, Hemostaseology, Infectious Diseases and Rheumatology, Oncologic Center, Paracelsus Medical University Salzburg

Daniel Egle, BrustGesundheitZentrum Tirol, Medical University Innsbruck

Mark Thill, National Coordinator, Agaplesion Markus Krankenhaus, Frankfurt

Jacqueline Sagasser, Klinikum Augsburg International Patient Service, Augsburg

Gerd Graffunder, Frauenarzt-Zentrum-Zehlendorf, Berlin

Dirk Behringer, Augusta-Kranken-Anstalt gGmbH Klinik, Bochum

Hans Tesch, Bethanien-Krankenhaus Onkologisches Zentrum, Frankfurt

Hans-Joachim Lück, Gynäkologisch-Onkologische Praxis, Hannover

Andreas Schneeweiss, NCT Nationales Centrum für Tumorerkrankunge, Universitätsklinikum Heidelberg, Heidelberg

Claudia Schumacher, Brustzentrum St. Elisabeth-Krankenhaus, Köln

Wolfram Malter, Uniklinik Köln Klinik und Poliklinic für Frauenheilkunde und Geburtshilfe Brestzentrum, Köln

Vladimir Semiglazov, National Coordinator, N.N. Petrov Research Inst. Oncol, St. Petersburg

Mona Frolova, N.N. Blokhin Medical Research Center of Oncology, Moscow

Alexander Vasiliev Gennadievich, Road clinical hospital of OJSC «Russian Railways», St.Petersburg

Nikita Volkov, St Petersburg Clinical Scientific Center, St. Petersburg

Begoña Bermejo, National Coordinator, Hospital Clinico Universitario, Valencia

Catalina Falo, Hospital Duran i Reynal Institut Català d’Oncologia, Hospitalet de Llobregat

Elena Sevillano, Hospital Universitario HM Sanchinarro, Madrid

Eva Maria Ciruelos Gil, Hospital Universitario 12 de Octubre, Madrid

José Ángel García Sáenz, Hospital Clínico San Carlos, Madrid

Anton Antón-Torres, Hospital Miguel Servet, Zaragoza

---

**Figure S1 – Association of cellularity with clinico-pathological features in the TCGA dataset** Pairwise differences in cellularity distribution between categorical variables were evaluated by Student’s *t* test. Association of cellularity with continuous variables was evaluated by Spearman’s correlation.

**Figure S2 – Association of cellularity with clinico-pathological features in the Park dataset** Pairwise differences in cellularity distribution between categorical variables were evaluated by Student’s *t* test. Association of cellularity with continuous variables was evaluated by Spearman’s correlation.

**Figure S3 – Association of cellularity with clinico-pathological features in thr Metzger-Filho dataset** Pairwise differences in cellularity distribution between categorical variables were evaluated by Student’s *t* test. Association of cellularity with continuous variables was evaluated by Spearman’s correlation.

**Figure S4 – Association of cellularity with clinico-pathological features in the NA-PHER2 dataset** Pairwise differences in cellularity distribution between categorical variables were evaluated by Student’s *t* test. Association of cellularity with continuous variables was evaluated by Spearman’s correlation.

**Figure S5 – Definition of candidate tumour-specific and stroma-specific genes across four breast cancer datasets a-c)** Single gene fold change values for one cancer specific (HALLMARK E2F TARGETS) and two stroma (HALLMARK EPITHELIAL MESENCHIMAL TRANSITION and Consensus TME^34^ T Cell CD8) specific genesets in the comparison between clinical specimens and matched patient-derived xenografts in the Bruna dataset. **d)** Distribution of correlation values between cellularity and 20844 genes in the NA-PHER2 dataset. Applied thresholds are indicated. **e)** Distribution of log fold change values for 16297 genes in the comparison between primary tumours and matched breast cancer PDXs. Applied thresholds are indicated. **f)** Distribution of correlation values between cellularity and 22189 genes in the Metzger-Filho dataset. Applied thresholds are indicated. **g)** Distribution of AUC values computed to quantify single gene expression ability to discriminate between medium/high and low/no tumour content in the NeoTRIP dataset. Applied thresholds are indicated.

**Figure S6 – Association of an immune score with cellularity and tumour purity signatures in the TCGA and Park datasets** Cellularity **(a,d)**, BCPS **(b,e)** and ESTIMATE score **(c,f)** were correlated with the immune score computed by Consensus TME^28^ in the TCGA **(a-c)** or Park **(d-f)** datasets (Spearman’s correlation).

## References

1. Hanahan D, Weinberg RA. Hallmarks of cancer: The next generation. Cell 2011;144(5):646–674; doi: 10.1016/J.CELL.2011.02.013/ATTACHMENT/3F528E16-8B3C-4D8D-8DE5-43E0C98D8475/MMC1.PDF.

2. Junttila MR, de Sauvage FJ. Influence of tumour micro-environment heterogeneity on therapeutic response. Nature 2013;501(7467):346–354; doi: 10.1038/NATURE12626.

3. Lou S, Zhang J, Yin X, et al. Comprehensive Characterization of Tumor Purity and Its Clinical Implications in Gastric Cancer. Front Cell Dev Biol 2022;9:3843; doi: 10.3389/FCELL.2021.782529/BIBTEX.

4. Zhang C, Cheng W, Ren X, et al. Tumor purity as an underlying key factor in glioma. Clinical Cancer Research 2017;23(20):6279–6291; doi: 10.1158/1078-0432.CCR-16-2598/116192/AM/TUMOR-PURITY-AS-AN-UNDERLYING-KEY-FACTOR-IN.

5. Aran D, Sirota M, Butte AJ. ARTICLE Systematic pan-cancer analysis of tumour purity. Nat Commun 2015;6; doi: 10.1038/ncomms9971.

6. Haider S, Tyekucheva S, Prandi D, et al. Systematic Assessment of Tumor Purity and Its Clinical Implications. JCO Precis Oncol 2020;4(4):995–1005; doi: 10.1200/PO.20.00016.

7. Fisher NC, Byrne RM, Leslie H, et al. Biological Misinterpretation of Transcriptional Signatures in Tumor Samples can Unknowingly Undermine Mechanistic Understanding and Faithful Alignment with Preclinical Data. Clinical Cancer Research 2022;28(18):OF1–OF14; doi: 10.1158/1078-0432.CCR-22-1102/706888/AM/BIOLOGICAL-MISINTERPRETATION-OF-TRANSCRIPTIONAL.

8. Rhee JK, Jung YC, Kim KR, et al. Impact of tumor purity on immune gene expression and clustering analyses across multiple cancer types. Cancer Immunol Res 2018;6(1):87–97; doi: 10.1158/2326-6066.CIR-17-0201/470743/AM/IMPACT-OF-TUMOR-PURITY-ON-IMMUNE-GENE-EXPRESSION.

9. Carter SL, Cibulskis K, Helman E, et al. Absolute quantification of somatic DNA alterations in human cancer. Nat Biotechnol 2012;30(5):413–421; doi: 10.1038/NBT.2203.

10. van Loo P, Nordgard SH, Lingjærde OC, et al. Allele-specific copy number analysis of tumors. Proc Natl Acad Sci U S A 2010;107(39):16910–16915; doi: 10.1073/PNAS.1009843107/-/DCSUPPLEMENTAL.

11. Bernard PS, Parker JS, Mullins M, et al. Supervised Risk Predictor of Breast Cancer Based on Intrinsic Subtypes. Journal of Clinical Oncology 2009;27(8):1160; doi: 10.1200/JCO.2008.18.1370.

12. Sparano JA, Gray RJ, Ravdin PM, et al. Clinical and Genomic Risk to Guide the Use of Adjuvant Therapy for Breast Cancer. N Engl J Med 2019;380(25):2395–2405; doi: 10.1056/NEJMOA1904819.

13. Callari M, Cappelletti V, D’Aiuto F, et al. Subtype-specific metagene-based prediction of outcome after neoadjuvant and adjuvant treatment in breast cancer. Clinical Cancer Research 2016;22(2):337–345; doi: 10.1158/1078-0432.CCR-15-0757/115760/AM/SUBTYPE-SPECIFIC-METAGENE-BASED-PREDICTION-OF.

14. Park YH, Lal S, Lee JE, et al. Chemotherapy induces dynamic immune responses in breast cancers that impact treatment outcome. Nat Commun 2020;11(1); doi: 10.1038/s41467-020-19933-0.

15. Smits AJJ, Kummer JA, de Bruin PC, et al. The estimation of tumor cell percentage for molecular testing by pathologists is not accurate. Modern Pathology 2014 27:2 2013;27(2):168–174; doi: 10.1038/modpathol.2013.134.

16. Anghel C v., Quon G, Haider S, et al. ISOpureR: an R implementation of a computational purification algorithm of mixed tumour profiles. BMC Bioinformatics 2015;16(1); doi: 10.1186/S12859-015-0597-X.

17. Ahn J, Yuan Y, Parmigiani G, et al. DeMix: deconvolution for mixed cancer transcriptomes using raw measured data. Bioinformatics 2013;29(15):1865–1871; doi: 10.1093/BIOINFORMATICS/BTT301.

18. Yoshihara K, Shahmoradgoli M, Martínez E, et al. Inferring tumour purity and stromal and immune cell admixture from expression data. Nature Communications 2013 4:1 2013;4(1):1–11; doi: 10.1038/ncomms3612.

19. Weinstein JN, Collisson EA, Mills GB, et al. The Cancer Genome Atlas Pan-Cancer Analysis Project. Nat Genet 2013;45(10):1113; doi: 10.1038/NG.2764.

20. Bruna A, Rueda OM, Greenwood W, et al. A Biobank of Breast Cancer Explants with Preserved Intra-tumor Heterogeneity to Screen Anticancer Compounds. Cell 2016;167(1):260–274.e22; doi: 10.1016/j.cell.2016.08.041.

21. Gianni L, Bisagni G, Colleoni M, et al. Neoadjuvant treatment with trastuzumab and pertuzumab plus palbociclib and fulvestrant in HER2-positive, ER-positive breast cancer (NA-PHER2): an exploratory, open-label, phase 2 study. Lancet Oncol 2018;19(2):249–256; doi: 10.1016/S1470-2045(18)30001-9.

22. Metzger-Filho O, Michiels S, Bertucci F, et al. Genomic grade adds prognostic value in invasive lobular carcinoma. Annals of Oncology 2013;24(2):377–384; doi: 10.1093/annonc/mds280.

23. Foroutan M, Bhuva DD, Lyu R, et al. Single sample scoring of molecular phenotypes. BMC Bioinformatics 2018;19(1):1–10; doi: 10.1186/S12859-018-2435-4/FIGURES/2.

24. Brueffer C, Vallon-Christersson J, Grabau D, et al. Clinical Value of RNA Sequencing-Based Classifiers for Prediction of the Five Conventional Breast Cancer Biomarkers: A Report From the Population-Based Multicenter Sweden Cancerome Analysis Network-Breast Initiative. JCO Precis Oncol 2018;2(2):1–18; doi: 10.1200/PO.17.00135.

25. Bianchini G, Pusztai L, Pienkowski T, et al. Immune modulation of pathologic complete response after neoadjuvant HER2-directed therapies in the NeoSphere trial. Ann Oncol 2015;26(12):2429–2436; doi: 10.1093/ANNONC/MDV395.

26. Gianni L, Huang CS, Egle D, et al. Pathologic complete response (pCR) to neoadjuvant treatment with or without atezolizumab in triple negative, early high-risk and locally advanced breast cancer. NeoTRIP Michelangelo randomized study. Annals of Oncology 2022; doi: 10.1016/j.annonc.2022.02.004.

27. Bianchini G, Qi Y, Alvarez RH, et al. Molecular anatomy of breast cancer stroma and its prognostic value in estrogen receptor-positive and -negative cancers. Journal of Clinical Oncology 2010;28(28):4316–4323; doi: 10.1200/JCO.2009.27.2419.

28. Jimenez-Sanchez A, Cast O, Miller ML. Comprehensive benchmarking and integration of tumor microenvironment cell estimation methods. Cancer Res 2019;79(24):6238–6246; doi: 10.1158/0008-5472.CAN-18-3560/653647/AM/COMPREHENSIVE-BENCHMARKING-AND-INTEGRATION-OF.

29. Symmans WF, Wei C, Gould R, et al. Long-Term Prognostic Risk After Neoadjuvant Chemotherapy Associated With Residual Cancer Burden and Breast Cancer Subtype. J Clin Oncol 2017;35(10):1049–1060; doi: 10.1200/JCO.2015.63.1010.

30. Quon G, Haider S, Deshwar AG, et al. Computational purification of individual tumor gene expression profiles leads to significant improvements in prognostic prediction. Genome Med 2013;5:29; doi: 10.1186/gm433.

31. Aran D, Sirota M, Butte AJ. Systematic pan-cancer analysis of tumour purity. Nat Commun 2015;6; doi: 10.1038/NCOMMS9971.

32. Symmans WF, Wei C, Gould R, et al. Long-Term Prognostic Risk After Neoadjuvant Chemotherapy Associated With Residual Cancer Burden and Breast Cancer Subtype. Journal of Clinical Oncology 2017;35(10):1049; doi: 10.1200/JCO.2015.63.1010.

33. Gianni L, Colleoni M, Bisagni G, et al. Effects of neoadjuvant trastuzumab, pertuzumab and palbociclib on Ki67 in HER2 and ER-positive breast cancer. NPJ Breast Cancer 2022;8(1); doi: 10.1038/s41523-021-00377-8.

34. Horr C, Buechler SA. Breast Cancer Consensus Subtypes: A system for subtyping breast cancer tumors based on gene expression. NPJ Breast Cancer 2021;7(1); doi: 10.1038/S41523-021-00345-2.

