## Supplementary figures and images for "Development and validation of a gene expression score to account for tumour purity and improve prognostication in breast cancer"

### Supplementary figures

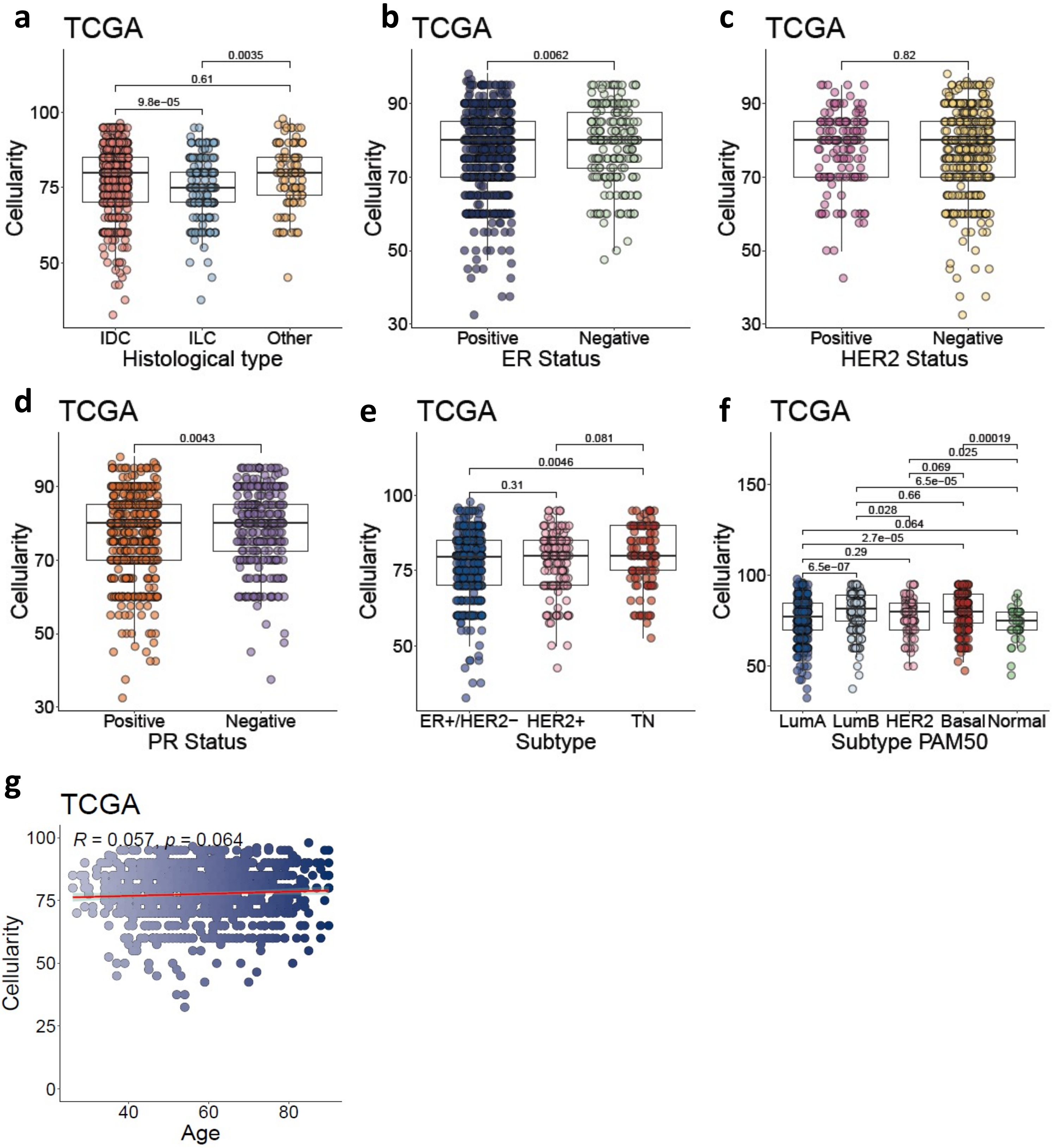

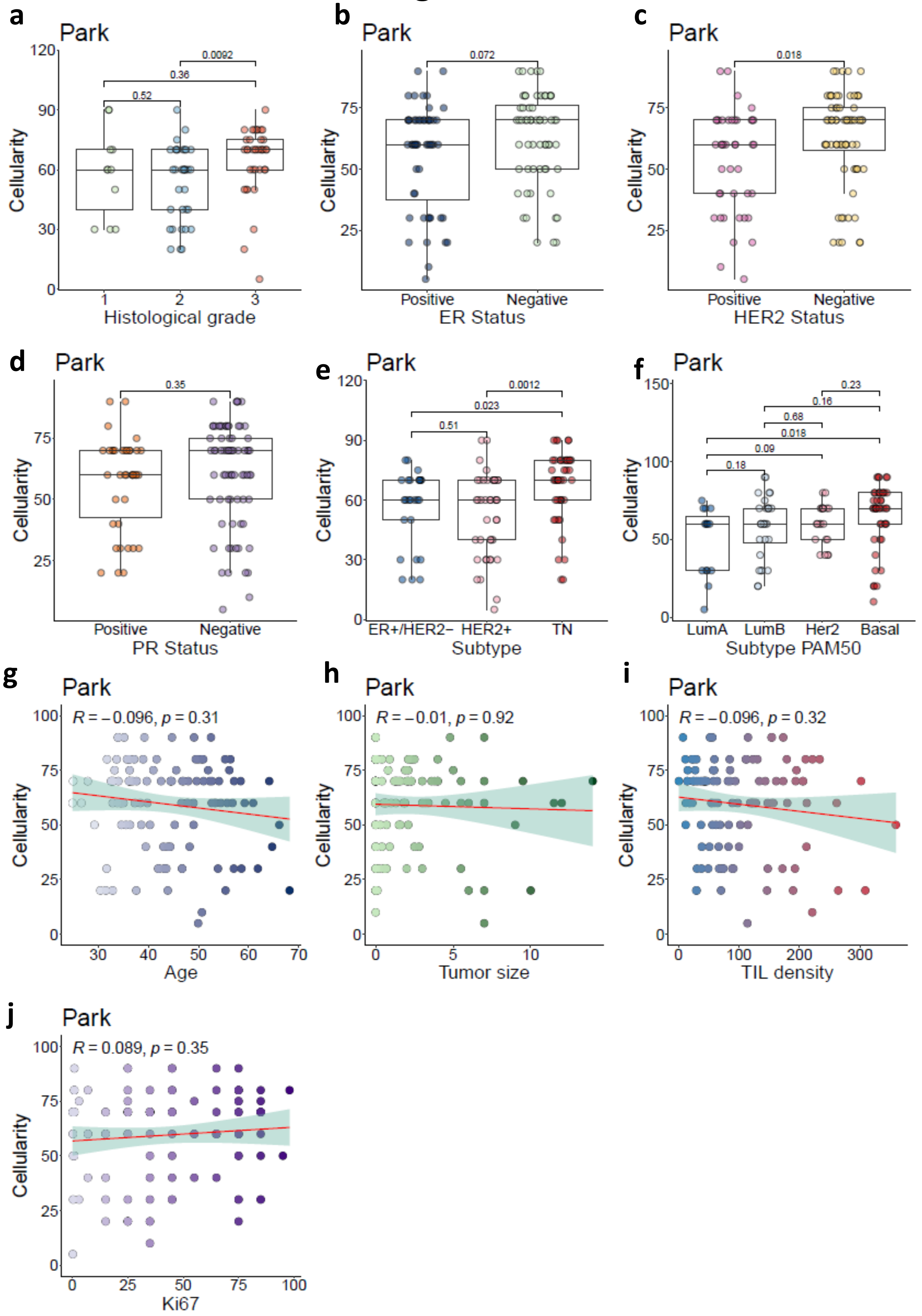

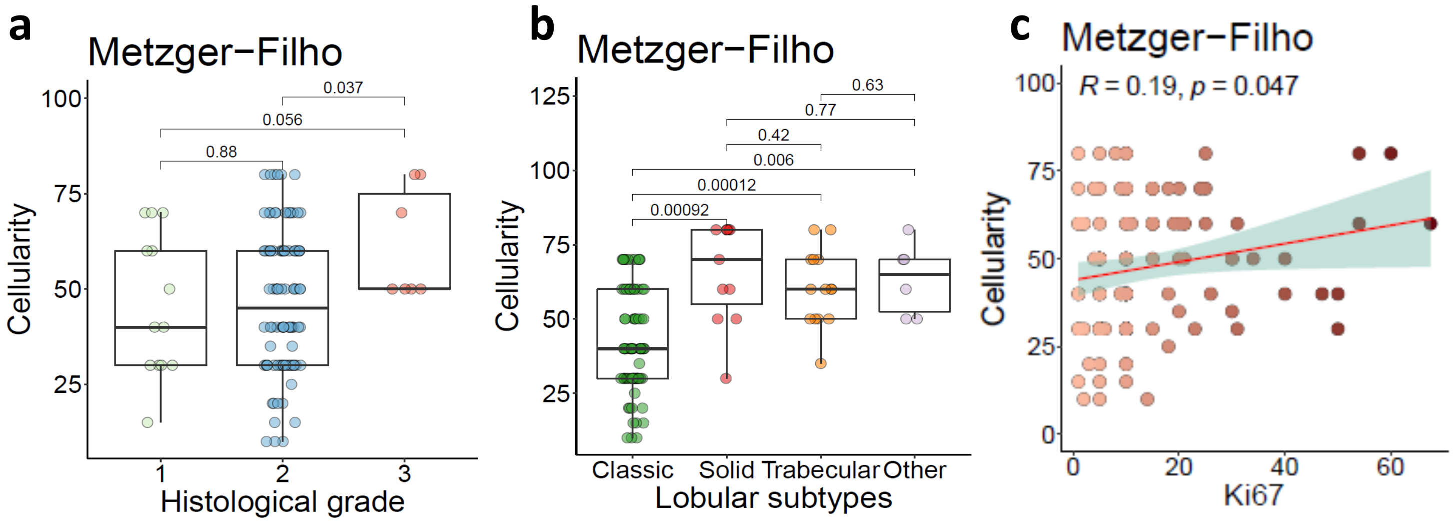

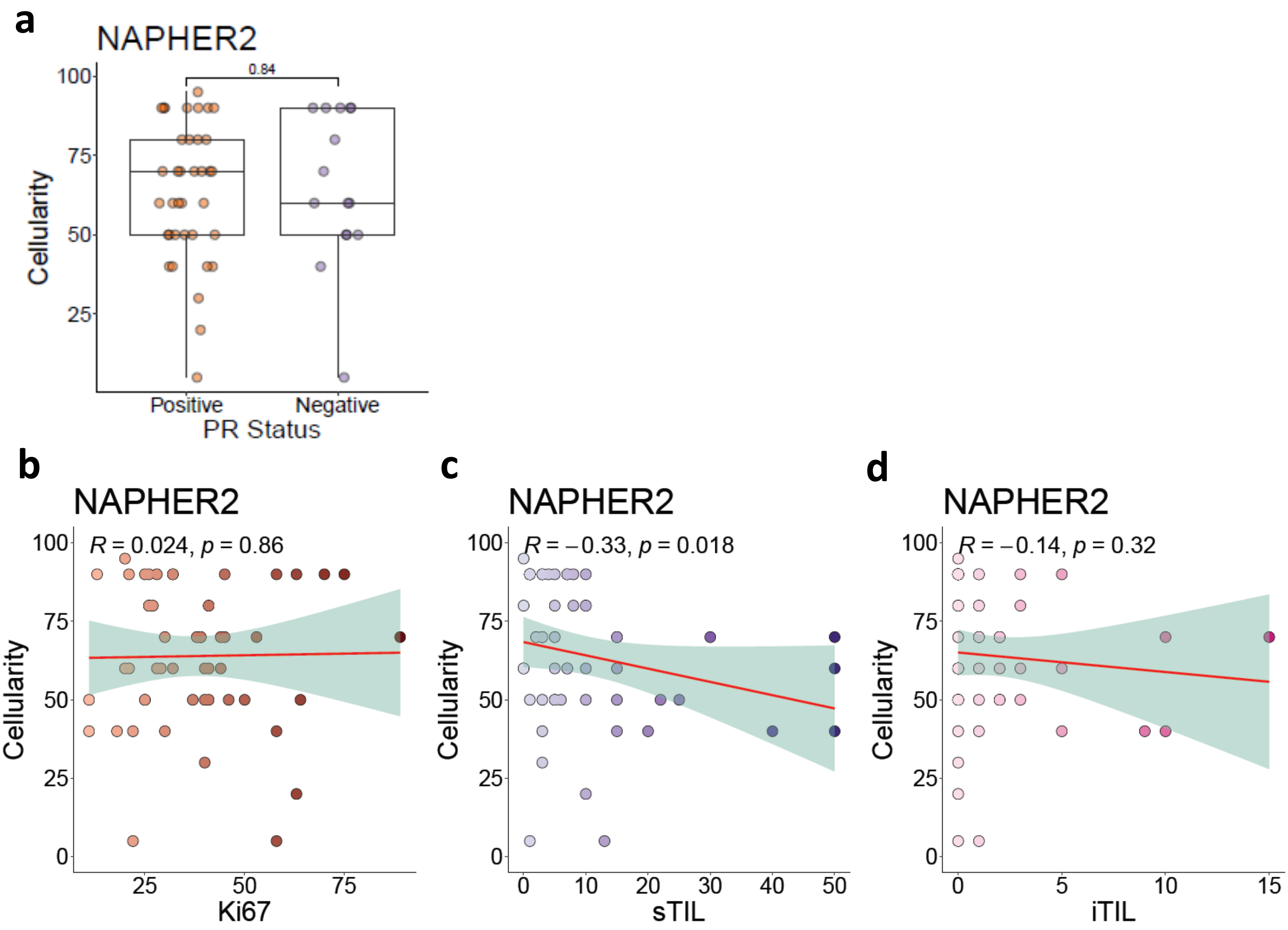

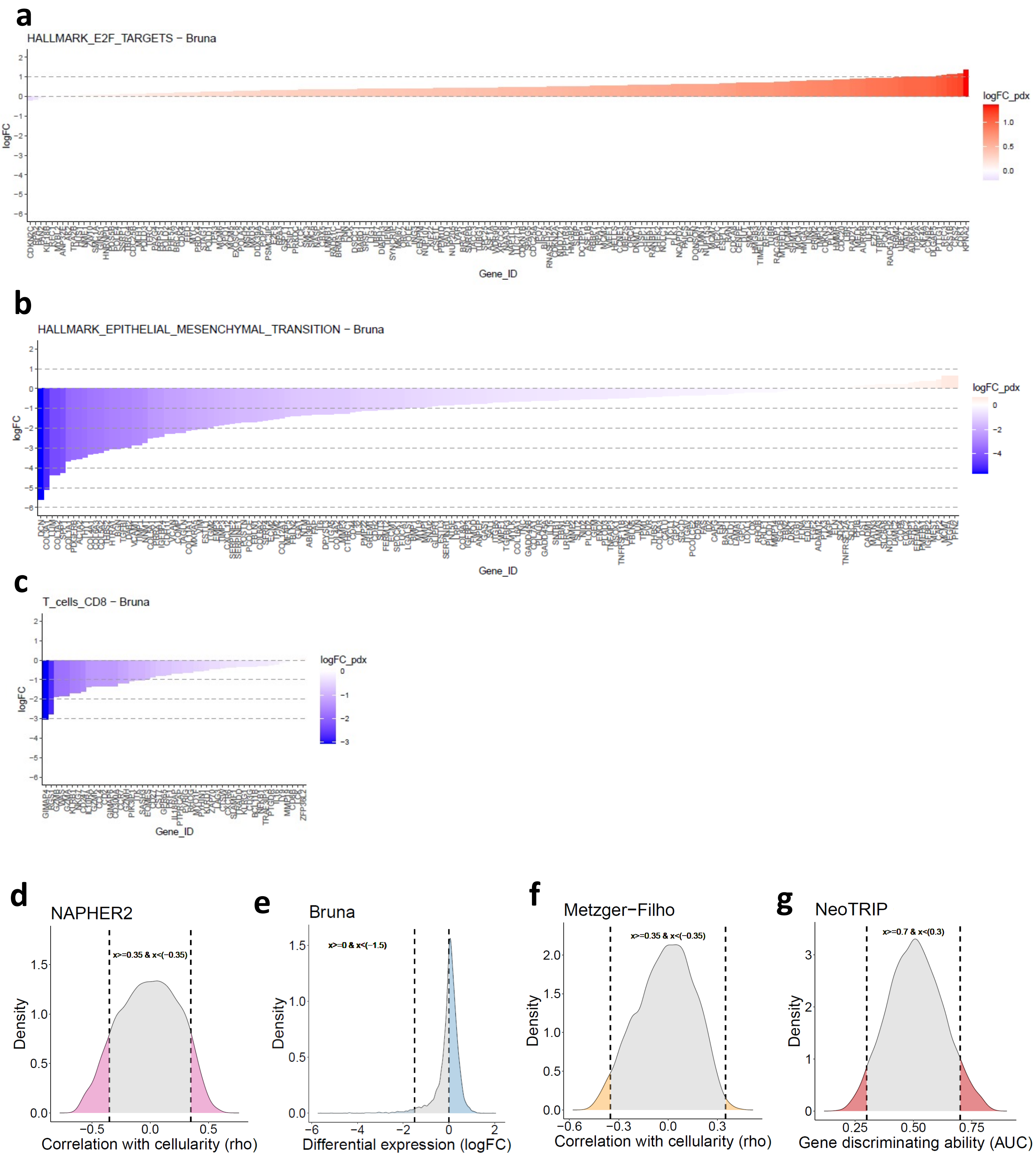

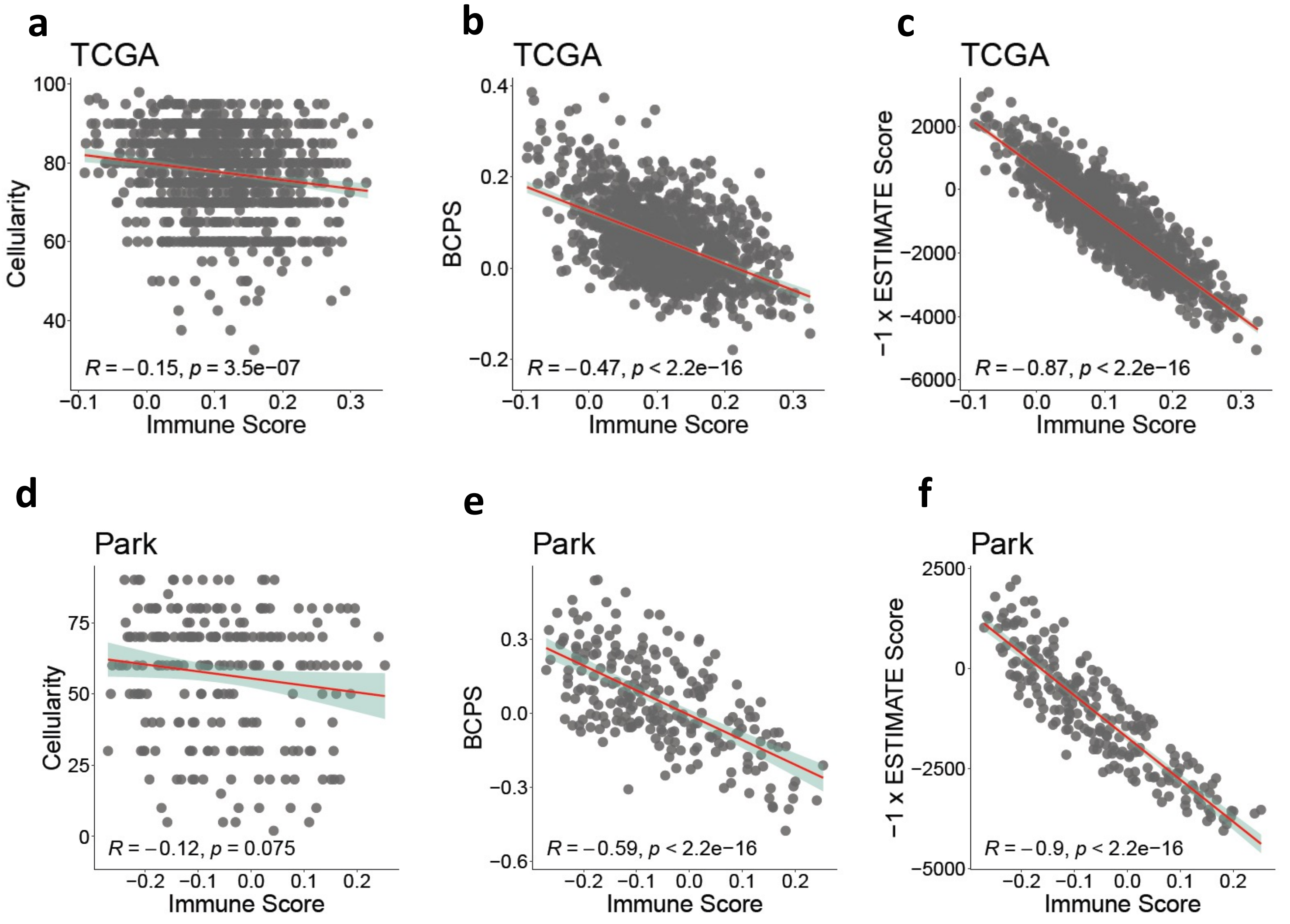
